# Genome-wide analysis of microRNA signature in lung adenocarcinoma with EGFR exon 19 deletion

**DOI:** 10.1101/032367

**Authors:** Lixia Ju, Mingquan Han, Xuefei Li, Chao Zhao

## Abstract

**Purpose:** The findings of EGFR mutations and the development of targeted therapies have significantly improved the overall survival of lung cancer patients. Still, the prognosis remains poor, so we need to know more about the genetic alterations in lung cancer. MicroRNAs are dysregulated in lung cancer, and microRNAs can regulate EGFR. So it is very important to predict the candidate microRNAs that target mutated EGFR and to investigate the role of these candidate microRNAs in lung cancer.

**Materials and methods:** In this study, we investigated the difference of microRNAs expression between lung adenocarcinoma cell lines with EGFR exon 19 deletion (H1650 and PC9) and its wild-type (H1299 and A549) using the Phalanx Human Whole Genome Microarray. Then the expression of individual microRNAs was validated by qRT-PCR assays. Moreover, we have detected the microRNAs expression in serum of lung adenocarcinoma patients with EGFR exon 19 deletion and wild-type.

**Results:** The expression of 1,732 microRNAs was evaluated, and we found that microRNAs expression was different between these two groups. Hsa-miR-141-3p, hsa-miR-200c-3p, hsa-miR-203, hsa-miR-3182, hsa-miR-934 were up-regulated and hsa-miR-3196 was down-regulated in the EGFR exon 19 deletion group compared with wild-type group. The detection of circulating microRNAs showed that miR-3196 was down-regulated in lung adenocarcinoma patients with EGFR exon 19 deletion compared with wild-type.

**Conclusions:** It is suggested that microRNAs associate with EGFR exon 19 deletion and miR-3196 can be further explored for potential predictor and targeted biomarker when it is difficult to get the tumors.

## Introduction

Lung cancer, one of the most common malignant tumor^1^, led to the most cancer death in the world. The prevalence and mortality of the lung cancer is still rising. In all new diagnosed lung cancer, non-small cell lung cancer (NSCLC) accounts for approximately 85%, which varies both in molecular and clinical presentation. Despite years of research, the 5-year survival is only about 18%.^2^ The discovery of key driver mutations has led to a more personalized approach in the treatment of advanced lung cancer. The activating mutations of EGFR implicate sensitivity to EGFR tyrosine kinase inhibitors. EGFR exon 19 deletion and L858R substitution in exon 21 have been extensively proved to be the sensitive mutations. However, the efficacy of EGFR-TKIs varies among different sensitive EGFR mutations. Several studies have reported that the advanced NSCLC patients with EGFR exon 19 deletion had a longer overall survival (OS) and/or progression-free survival (PFS) with the treatment of gefitinib or erlotinib compared with those with the L858R mutation. ^3–6^

MicroRNAs are evolutionarily conserved, endogenous small non-coding RNAs with 18-25 nucleotides that have important functions in diverse biological processes, such as cell proliferation, differentiation, and apoptosis.^7–9^ Furthermore, microRNAs play an essential role in behaving either as oncogenes or tumor suppressor genes^10^. Increasing evidence indicates that dysregulation of specific microRNAs contributes to the development and progression of cancer, including lung cancer.^11–13^ Moreover, microRNAs can regulate EGFR.^14^ So it is important to make it clear that the candidate microRNAs regulating EGFR exon 19 deletion because it might spark the design of novel therapeutics to combat the resistance to EGFR-TKIs or develop new targeted therapy.

In this study, we conducted an explorative microRNAs expression study in two groups of lung adenocarcinoma cell lines, including EGFR exon 19 deletion group (H1650 and PC-9) and EGFR wild-type group (A549 and H1299), using microRNA microarrays. Our main focus was the different microRNAs in the two groups. The selected microRNAs were confirmed using qRT-PCR in the cell lines and then we measured 3 microRNAs expressed differently in cell lines in the serum of 14 non-smoking female lung adenocarcinoma patients with wt EGFR and 13 patients with EGFR 19 del. Accordingly we discussed the association of microRNAs with EGFR exon 19 deletion.

## Material and Methods

### Cell lines and cell culture

Human adenocarcinoma cell lines PC-9 and H1650 (harboring EGFR exon 19 deletion) and A549 and H1299 (harboring wild-type EGFR) were provided by Cancer Institute of Tongji University Medical School, China. All these cells were cultured at 37°C with 5% CO2 in Dulbecco’s modified Eagle’s medium (DMEM) supplemented with 10% fetal bovine serum (FBS), 100 U/ml penicillin and 100 mg/ml streptomycin.

### Patient enrollment and serum samples

All patients in the study were recruited form Shanghai Pulmonary Hospital, Tongji University Medical School, between January 2012 and June 2014. The patients were newly diagnosed and histologically confirmed primary lung adenocarcinoma. Patients with the previous history of cancer, radiotherapy or chemotherapy were excluded. The study was approved by an ethical review committee at Tongji University Institutional Care and Use Committee. All blood serum samples were collected and put into a liquid nitrogen tank for long-term storage until microRNAs extraction.

### Total RNA isolation

Total RNA was extracted from cells using TRIZOL Reagent (Invitrogen, USA). The RNA concentration and purity were accessed by OD260/OD280 (≥1.6) and OD260/OD230 (≥1.0), and the RNA yield and quality were checked (RIN≥5.0) using Agilent 2100 Bioanalyzer (Agilent Technologies, Santa Clara, CA, USA).

### Human microRNAs OneArray®

Human microRNA OneArray® v3 (Phalanx Biotech Group, Taiwan) contains triplicated 1,711 unique microRNA probes from Human (miRBase Release v17), each printed in technical triplicate, and 189 experimental control probes.

### Microarray analysis

Small RNA was pre-enriched by Nanoseplook (Pall Corporation, USA) from 2.5 μg total RNA samples and labeled with microRNAs ULSTM Labeling Kit (Kreatech Diagnostics, The Netherlands). Labeled targets were hybridized to the Human microRNA OneArray® v3 with OneArray® Hybridization System. After 16 hours hybridization at 37 □, non-specific binding targets were washed away by three different washing steps (Wash □ 37 □ 5 mins; Wash □37 □, 5 mins 25 □ 5 mins; Wash III rinse 20 times), and the slides were dried by centrifugation and scanned by an Axon 4000B scanner (Molecular Devices, Sunnyvale, CA, USA). Normalized spot intensities were transformed to gene expression log2 ratios between the mutation and wild-type group. The spots with log2 ratio ≥ 1 or log2 ratio ≤ -1 and P-value 6 0.05 were tested for further analysis.

### Validation by qRT-PCR

Quantitative real time-PCR (qRT-PCR) was carried out on the 7900HT thermocycler (Applied Biosystems, Foster City, CA). U6 was used for internal controls. The data were managed using the Applied Biosystems software RQ. Relative expression was calculated using the comparative Ct method and obtaining the fold-change value (2^−ΔΔCt^). Data analyses were performed via GraphPad Prism v6.00.

### EGFR mutation analyses

Mutation analyses of EGFR exons 18-21 were performed on 27 of the tumor samples using the amplification-refractory mutation system assay (ARMS). Data analyses were performed by employing the LightCycler Adapt software (LightCycler 480 Software, v. 1.5).

### Digital PCR

Digital PCR was performed in parallel for the measurement of microRNAs in the serially-diluted oligonucleotides. 30 μL of the reaction mixture containing 15 μL QuantStudio^TM^ 3D Digital PCR Master Mix, 2X (Life technologies), 2 μL of cDNA solution, and 1.5 μL of TaqManR Assay, 20X (primer/probe mix) (Life technologies) and 11.5 μL water. The droplets generated from each sample were transferred to a 96-well PCR plate (Eppendorf, Germany). PCR amplification was carried on a T100 thermal cycler (QuantStudio^TM^ 3D Digital PCR System) at 96°C for 10 min, followed by 39 cycles 60°C for 2 min, then 98°C for 0.5 min. For Final extension followed by 60°C for 2 min, Last, Storage is at 10°C for 100min.^15^

## Results

### Microarrays analysis of cell lines with EGFR exon 19 deletion compared with wild-type

To identify microRNAs that were differentially expressed between cell lines with EGFR exon 19 deletion and the wild-type adenocarcinoma cell lines, the expression profiles of microRNAs (1,711 microRNAs) were assessed using microRNA microarrays. For clustering analysis, 260 genes was selected to identify differentially expressed based on the threshold of fold change and p-value (Figure 1). Standard selection criteria to identify differentially expressed genes are established at log2 |Fold change | ≥ 0.8 and P < 0.05. The microRNAs differentially expressed between EGFR exon 19 deletion and wild-type are shown in Table 1, (*P* < 0.05). In total, let-7d-3p, miR-1307-5p, miR-141-3p, miR-200c-3p, miR-203, miR-3182, miR-4510, miR-934 were up-regulated in EGFR exon 19 deletion group compared with the wild-type group. On the contrary, miR-3196, miR-4450, miR-4649-5p were down-regulated.

**Figure 1.**
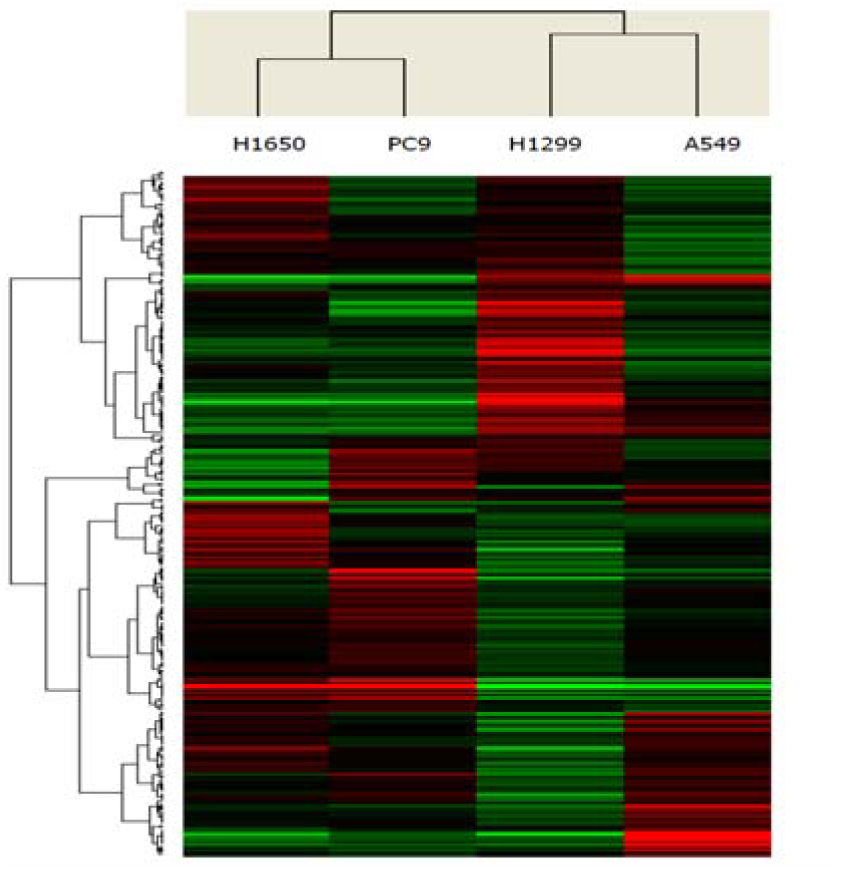
Unsupervised clustering of 260 microRNAs (rows) in 4 lung adenocarcinoma cell lines (columns). Up- and down-regulated genes are represented in red and green colors, respectively.

**Table 1.**
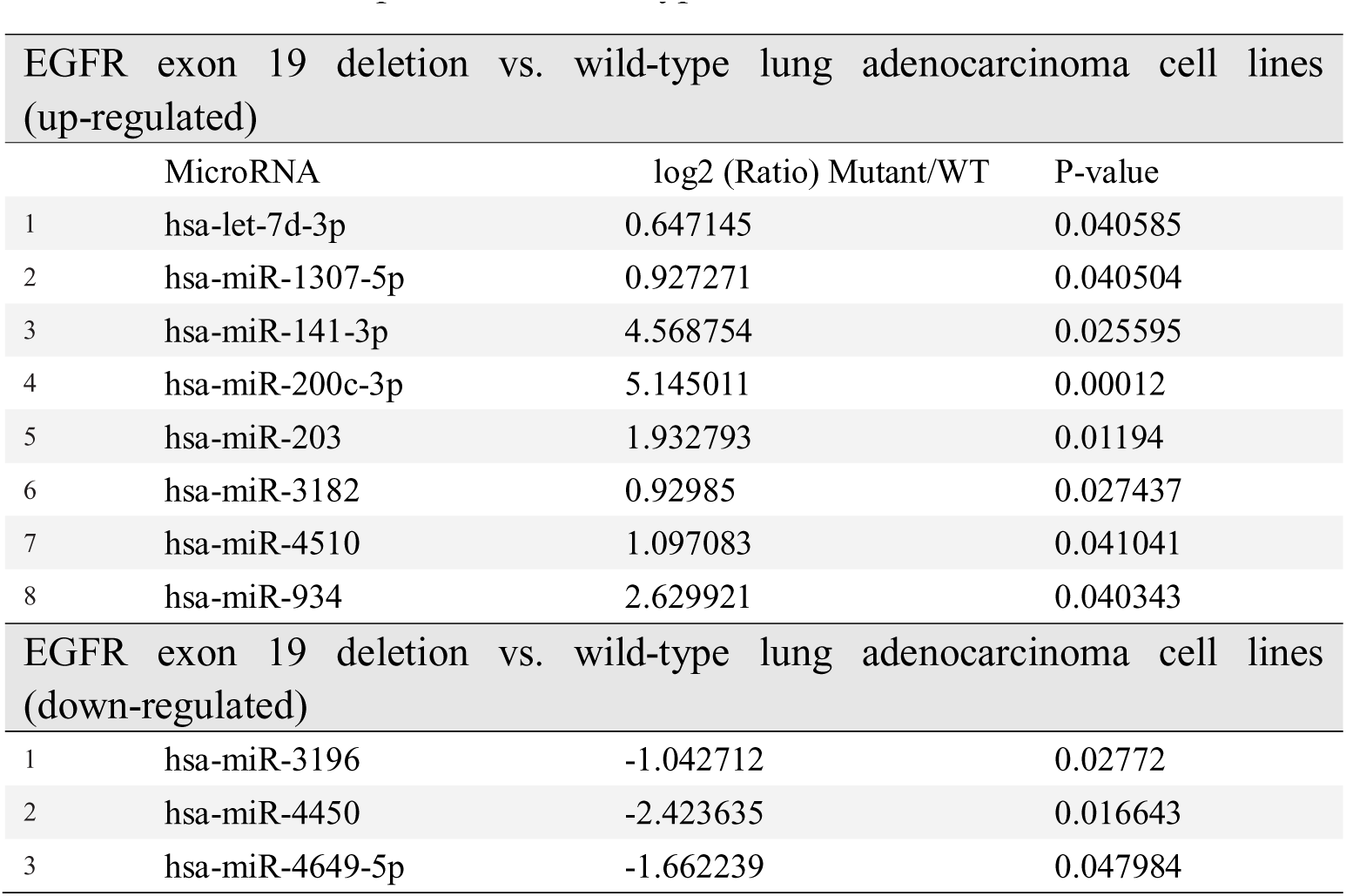
MicroRNAs that were significantly differentially expressed between EGFR exon 19 deletion compared with wild-type cell lines

### Validation of the microarrays results using quantitative reverse transcriptase polymerase chain reaction (qRT-PCR)

The eleven microRNAs, differentially expressed in the two cell groups in microarray analyses, were validated by qRT-PCR. As a result, six microRNAs were identified and differentially expressed between the two cell groups (see Table 2). Hsa-miR-141-3p, hsa-miR-200c-3p, hsa-miR-203, hsa-miR-3182, and hsa-miR-934 were up-regulated, while hsa-miR-3196 was down-regulated (Figure 2).

**Table 2.**
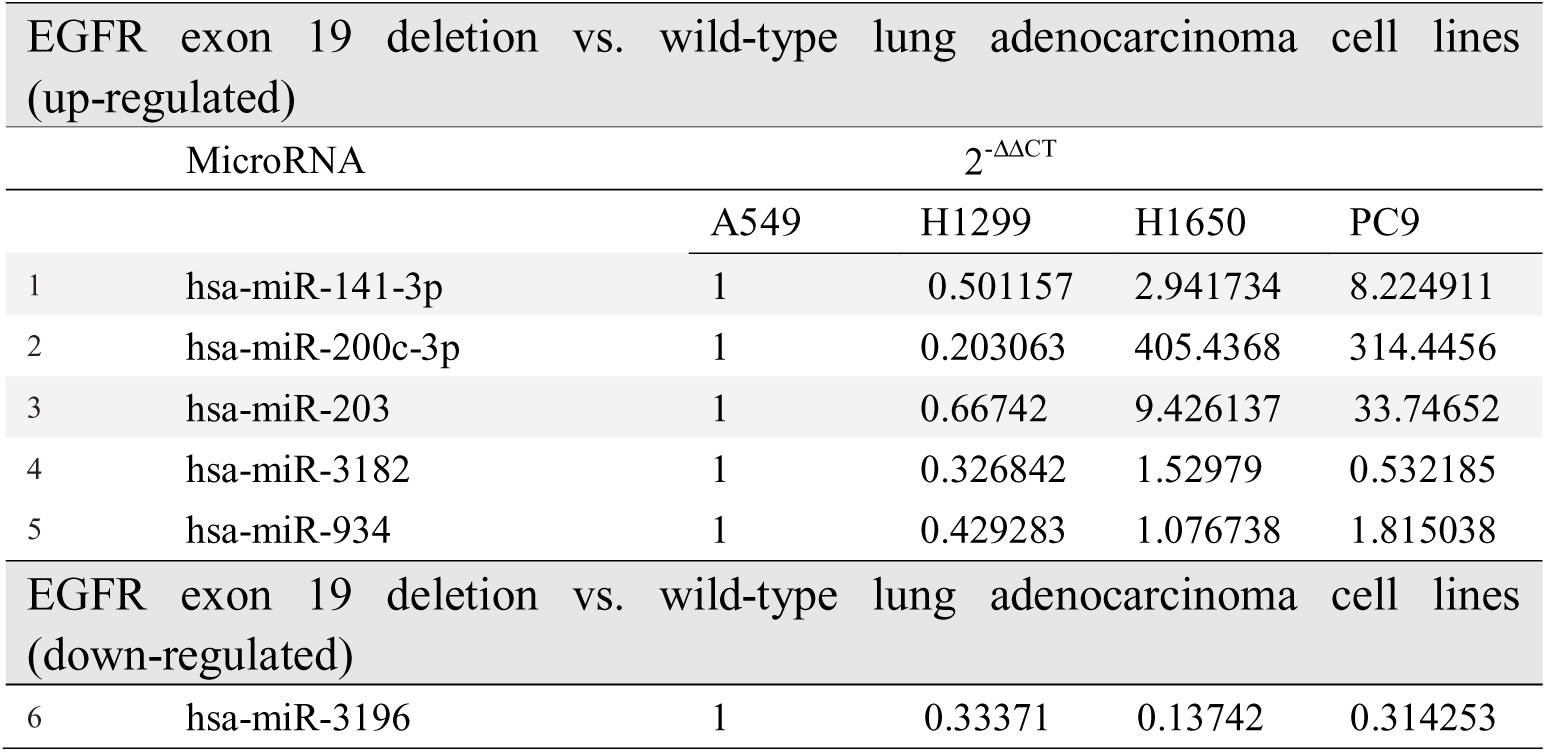
Validation of the microRNAs that were significantly differentially expressed between EGFR exon 19 deletion compared with wild-type cell lines (qRT-PCR)

**Figure 2.**
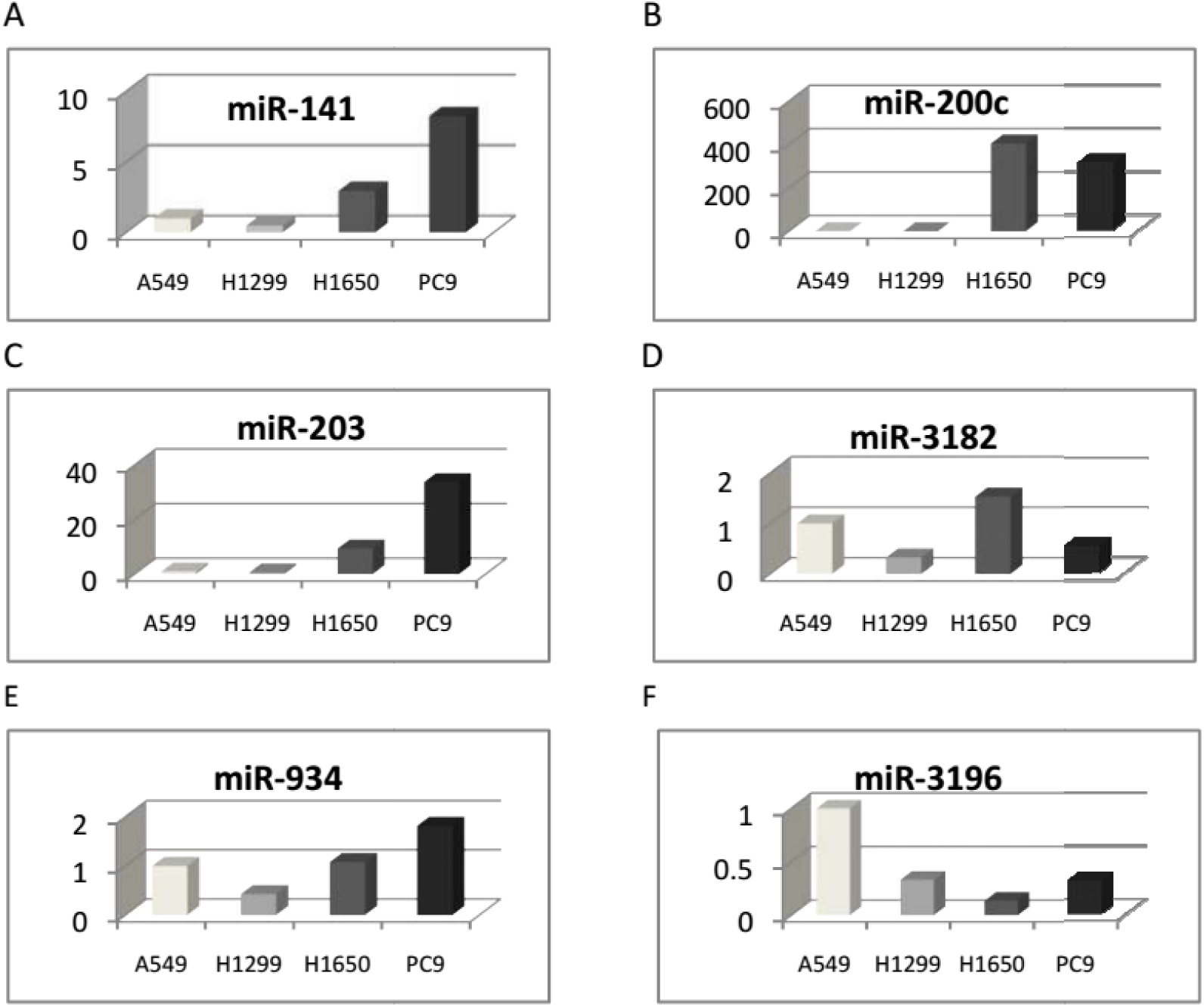
Six microRNAs were identified and differentially expressed between the two cell groups with EGFR exon 19 deletion and wild-type using qRT-PCR. A: Hsa-miR-141-3p, B: hsa-miR-200c-3p, C: hsa-miR-203, D: hsa-miR-3182, E: hsa-miR-934 was up-regulated, and F: hsa-miR-3196 was down-regulated.

### Pathway analysis

Using the microRNA target filter in TargetScan Release 6.2, we searched for genes involved in the signaling pathways that are experimentally observed or highly predicted to be regulated by the selected microRNAs. The six microRNAs that were differentially expressed between EGFR exon 19 deletion and EGFR wild-type cell lines were associated with 3181 mRNA targets. Two microRNAs (hsa-miR-200c-3p, hsa-miR-203) were experimentally confirmed to target EGFR, and be associated with EGFR-TKI resistance.^16–17^

### Circulating microRNAs in relation to EGFR exon 19 deletion

This study was conducted on 27 participants stratified into EGFR exon 19 deletion group and wild-type group. Selected candidates were non-smoking lung adenocarcinoma patients. It is well known that the sensitivity of qPCR for the detection of the low copy genes is not so high, as it only can resolve ~1.5-fold changes of nucleic acids.^18^ Given that a portion of the cancer-related microRNAs is derived from primary tumor and could be ‘diluted’ in the normal microRNAs,^19–21^ the microRNAs presenting at low levels in serum could be undetectable by qPCR.Droplet digital PCR is a direct method for quantitatively detecting nucleic acids.^22,23^ It depends on limit ptoportition of the PCR volume, in which a positive microreactions indicates the presence of a single molecule in a given reaction. The number of positive reactions, together with Poisson’s distribution, can be used to calculate the original target concentration in a straight and high-confidence measurement method.^24^ In our study, digital PCR was used to detect three circulating microRNAs (hsa-miR-200c-3p, hsa-miR-203, hsa-miR-3196) of 27 patients, 13 patients with EGFR exon 19 deletion and 14 wild-type EGFR. In the results, we found that hsa-miR-3196 was down-regulated in the group of EGFR exon 19 deletion compared with wild-type, which was consistent with our results of cell study (Figure 3).

**Figure 3.**
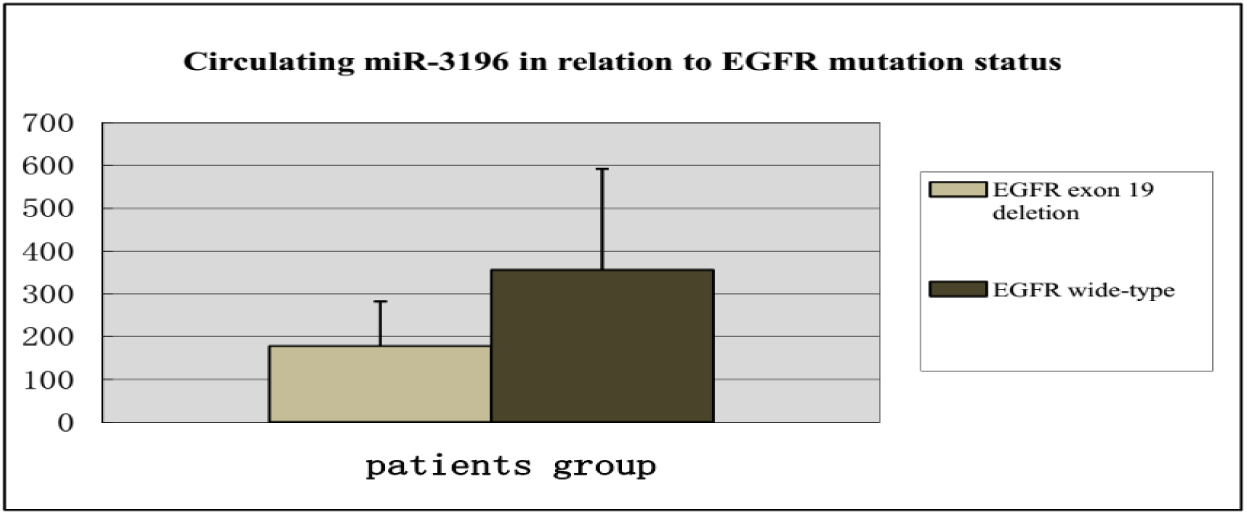
Digital PCR results showed that circulating hsa-miR-3196 was down-regulated in serum of lung adenocarcinoma patients with EGFR exon 19 deletion compared with wild-type.

## Discussion

In the present work, we found eleven microRNAs differentially expressed between EGFR exon 19 deletion cell lines and EGFR wild-type lung adenocarcinoma cell lines in the microarray analysis. Then the PCR study identified that hsa-miR-141-3p, hsa-miR-200c-3p, hsa-miR-203, hsa-miR-3182, and hsa-miR-934 were up-regulated, and hsa-miR-3196 was down-regulated in EGFR exon 19 deletion group compared with wild-type group. Moreover, the detection of circulating microRNAs found that miR-3196 was down-regulated in patients with EGFR exon 19 deletion compared with wild-type lung adenocarcinoma patients.

Presently, not much is known about the microRNAs expression in EGFR exon 19 deletion versus wild-type lung adenocarcinomas. Using microRNA microarrays,Dacic et al. analyzed six lung adenocarcinomas and found that miR-155 was up-regulated only in EGFR/KRAS-negative group, miR-25 was up-regulated only in EGFR-positive group and miR-495 was up-regulated only in KRAS-positive adenocarcinoma. In opposite, let-7g was down-regulated in all three groups, with more significant downregulation in EGFR/KRAS-negative adenocarcinoma.^25^ Seike et al. identified 12 microRNAs differentially expressed between 6 EGFR-mutated tumors and 22 wild-type tumors. Zhang et al. found that miR-122 were differentially expressed between wild and mutant EGFR carriers (P=0.018).^26^ But all these have not accurately make the relation of microRNAs and EGFR exon 19 deletion clear.

In our results, Hsa-miR-141-3p and hsa-miR-200c-3p were up-regulated in exon 19 deletion versus wild-type. MiR-141-3p is a miR-200 family member, which consists of five microRNAs located in two different clusters (miRs-200b/a/429 and miRs-200c/141) on chromosome 1 and 12 in humans.^27^ Previous studies have shown that miR-141/miR-200 is involved in cancer development and metastasis.

Tejero R et al. reported that high miR-141 and miR-200c expression are associated with shorter OS in NSCLC patients with adenocarcinoma through MET and angiogenesis.^28^ MiR-141 and miR-200c expression was significantly up-regulated in NSCLC tissues, and its overexpression accelerated NSCLC cell proliferation in vitro and tumor growth in vivo.^29,30^ In line with our results, miR-203 overexpression resulted in increased sensitivity to gefitinib-induced apoptosis in nude mice after two weeks of treatment.^31^

The function of hsa-miR-934 is unknown, but it is located in intron 4 of the vestigial-like 1 (VGLL1) gene. The miR-934 was the most strongly up-regulated microRNA in triple-negative IDCs (61.5-fold increase with respect to ER+ breast carcinomas).^32^ Though studies of hsa-miR-3196 are rare, it was found that miR-3196 was down-regulated in basal cell carcinoma compared with nonlesional skin,^33^ and was also down-regulated in PTC patients with non-(131) I-avid lung metastases versus (131)I-avid lung metastases.^34^ Regarding hsa-miR-3182, there is still no published study.

A large number of microRNAs have been found to be stably expressed in human serum and plasma^35,36^. Circulating microRNAs or their expression profiles have been proposed to be useful biomarkers of the diagnosis and prognosis of cancer. Overexpressed serum miR-21 was associated with the poor survival, lymph node metastasis and advanced stage of NSCLC.^37^

As previously stated, very little is known about miR-3196, but the data from the next two previous published studies point to the possibility that EGFR exon 19 deletion could have a distinct molecular identity. Bjaanaes MM et.al examined microRNA expression in 154 surgically resected lung adenocarcinomas and 20 corresponding normal lung tissue samples using Agilent microarrays and found that 17 microRNAs were differentially expressed between EGFR-mutated and EGFR wildtype tumors.^38^ Recently, Zhang et al. reported that circulating miR-195 and miR-122 may play important roles in predicting the overall survival as well as predicting EGFR mutation status in non-smoking female patients with lung adenocarcinoma. Measuring plasma levels of miR-195 and miR-122 may have especial values for EGFR mutant patients with lung adenocarcinoma.^39^ In this regard, we selected hsa-miR-200c-3p, hsa-miR-203, and hsa-miR-3196 to measure the expression level in serum of patients. The reason of our selection was the high expression level in cell lines. Lastly, hsa-miR-3196 was significantly differentially expressed between EGFR exon 19 deletion compared with wild-type lung adenocarcinomas. It is consistent with our result of cell lines.

Therefore, the circulating microRNAs may potentially provide a noninvasive strategy for predicting response to EGFR TKIs when it’s difficult to get the tumor tissues. Moreover, we identified that microRNAs regulate EGFR exon 19 deletion and which may represent a clinically useful modality to treat TKI resistance in NCSLC patients.

## Acknowledgements

This work was supported by The National Natural Science Foundation of China (81207106) and Shanghai Municipal Health Bureau Fund (20124Y123; 2012L051A); and the Program of Science and Technology Commission of Shanghai Municipality (14401932200; 12401907500). We also thank Prof. Rongzheng, Ren for his revision of the draft.

